# DevKidCC allows for robust classification and direct comparisons of kidney organoid datasets

**DOI:** 10.1101/2021.01.20.427346

**Authors:** Sean B. Wilson, Sara E. Howden, Jessica M. Vanslambrouck, Aude Dorison, Jose Alquicira-Hernandez, Joseph E. Powell, Melissa H. Little

## Abstract

Kidney organoids provide a valuable resource to understand kidney development and disease. Clustering algorithms and marker genes fail to accurately and robustly classify cellular identity between human pluripotent stem cell (hPSC)-derived organoid datasets. Here we present a new method able to accurately classify kidney cell subtypes, a hierarchical machine learning model trained using comprehensive reference data from single cell RNA-sequencing of human fetal kidney (HFK). We demonstrate the tool’s (*DevKidCC*) performance by application to all published kidney organoid datasets and a novel dataset. *DevKidCC* is available on Github and can be used on any kidney single cell RNA-sequence data.

## Background

Single cell RNA sequencing has reformed how we approach biological questions at the transcriptional level, facilitating accurate evaluation of cellular heterogeneity within complex samples, including entire tissues. When coupled with approaches for molecular lineage tagging^1^ and computational approaches to analyse pseudotime^2–4^ and RNA velocity^5,6^, gene expression in complex tissues such as kidney can be studied at an unprecedented resolution. Despite these advantages, classification of cellular identity remains challenging and variable between data, even when analysing similar cellular systems. Currently, a common approach for identifying cell populations within single cell data is to first cluster cells, compute differentially expressed genes between clusters, and label clusters of cells based on expression of known marker genes^4,7,8^. The choice of clusters can be arbitrary, with users defining the number of clusters, thereby raising the potential for biases in the reproducibility of cell-type labels^9^. Placement of cells into a cluster relies on transcriptional similarity^10^, hence there needs to be a large enough population with a distinct gene signature for this to occur. Cell clusters are also commonly defined based upon one or a few known differentially expressed genes rather than their global transcriptional signature. Finally, technical challenges such as batch variation can impact definitive cellular identification.

The application of single cell profiling to developmental biology presents unique challenges due to the presence of intermediate cell types undergoing differentiation during morphogenesis. The mammalian kidney contains more than 25 cell types in the mature postnatal tissue, arising from a smaller number of progenitor cell types including nephron, stromal, endothelial and collecting duct progenitors. Organogenesis is driven via reciprocal signalling and self-organisation with many intermediate transcriptional states that are less well defined, making the classification of cell types at the single cell level both extremely useful but particularly difficult (reviewed in Little and Combes, 2019^11^). This is further complicated with hPSC-derived kidney organoid datasets. While protocols for differentiating kidney organoids from hPSC attempt to replicate *in vivo* kidney differentiation, they are likely to be limited and contain emerging non-specific, off-target or synthetic cell types^12–15^. Here, unbiased classification of cellular identity is a computational challenge. Indeed, recent single cell profiling of cell human fetal kidney (HFK) datasets have shown that the classical canonical markers for many cell identities within the kidney are not unique to these cell types but are also expressed at lower levels within other populations^15–18^. This makes cell classification in organoids more challenging when analysing gene expression of these markers in the single cell clusters. The ability to robustly identify and classify cells in hPSC-derived organoid data is crucial to facilitate useful comparisons between datasets, particularly data generated using different differentiation protocols and cell lines but also in response to mutation or perturbation. These analyses will also help to improve and refine protocols towards a more accurate endpoint tissue.

One approach to cellular identification is to apply a small set of ‘known’ genes to identify clusters within a dataset based upon an existing reference dataset that has been accurately classified. Of the 12 kidney organoid single cell RNA-seq datasets published to date (Table 1), seven used a HFK reference to find congruence with their clustered organoid populations either through integration or training a unique random forest classifier. However, there have been many different references used across these publications. Cell classifications may be inconsistent when using various references containing different proportions of cells, possibly captured at different ages or regions of the tissue. Indeed, the most commonly used HFK reference only contained cells from the cortex of a 16-week kidney and hence was reported to contain few nephron cells and no ureteric epithelium^19^. There have been many tools developed to utilise reference data to classify a related query dataset, with scrna-tools.org^4^ listing 85 tools in the “Classification” category. These tools extract cell type information from an annotated reference and apply that to a query dataset. Most rely upon the user to supply the reference data and for those that supply a reference, none are directly relevant to hPSC-derived kidney organoids. The *R* packages *scTyper*^20^ and *scClassify*^21^ are trained on existing datasets. These are not ideal for human developing kidney classification as *scClassify* is trained on mouse cell data, while *scTyper* contains gene sets of limited cell types of the adult kidney and is thus not a developing kidney cell population. As such there is a no tool that can be used to directly and accurately classify the cell types present within the developing human kidney.

**Table 1:** Summary of existing kidney related single cell datasets

| Human Fetal Kidney |  |  |  |  |
| --- | --- | --- | --- | --- |
| Reference | Age (post coitum) | Sample Details |  |  |
| Lindstrom *et al.*<sup>19</sup> | 16 weeks | MARIS dissociation used to isolate cortical regions |  |  |
| Menon *et al.*<sup>28</sup> | 87-132 days | Cells from cortical nephgenic zone to inner medullary region |  |  |
| Young *et al.*<sup>29</sup> | 1.9 and 2.1 months | Biopsies of 1.9 and 2.1 month fetal kidneys |  |  |
| Hochane *et al.*<sup>25</sup> | 9, 11, 13, 15, 18 weeks | Week 9, 11, 13, 16 and 18 kidney pieces |  |  |
| Tran *et al.*<sup>26</sup> | 15 and 17 weeks | Regions dissected from both inner and outer cortex |  |  |
| Holloway *et al.*<sup>23</sup> | 16 weeks | Wedge biopsy including both medulla and cortex, one day 96 male and one day 108 female sample |  |  |
| Kidney Organoids |  |  |  |  |
| Reference | Age (days) | Sample information | ID | Classification |
| Wu *et al.*<sup>12</sup> | 26 | 4 batches of iPS and 2 batches of ES derived organoids using Takasato<sup>30</sup> protocol | Wu_T | Clustering & DE genes, Integration with self-generated adult snRNA dataset, Lindstrom<sup>19</sup> trained random forest classifier |
|  | 26 | 3 batches of iPS and 1 batch of ES derived organoids using Morizane<sup>31</sup> protocol | Wu_M |  |
|  | 34 | Older iPS derived organoid using Takasato protocol | Wu_TO |  |
|  | 7, 12, 19, 26 | Time course of iPS derived organoids using Takasato protocol | Wu_TC |  |
|  | 26 | 2 batches of iPS derived Takasato organoids with BDNF inhibition | Wu_TB |  |
| Czerniecki *et al.*<sup>32</sup> | 25 | Freedman iPS and ES derived organoids, modified protocol for High Throughput Sequencing, comparing with/without VEGF | Cz_F | Clustering & DE genes, Menon |
| Howden *et al.*<sup>13</sup> | 18, 25 | Takasato iPS derived organoids using E6 base media | How_T | Clustering & DE genes |
| Phipson *et al.*<sup>33</sup>, Combes *et al.*<sup>15</sup> | 25 | Takasato iPS derived organoids generated in two batches. Same dataset in both publications | PC_T | Clustering & DE genes; Integration with Lindstrom<sup>19</sup> |
| Harder *et al.*<sup>34</sup> | 19 | Freedman ES derived organoids, 6 datasets generated from the all organoids in a well, 3 separate batches | Har_F | Clustering & DE genes, integration and trajectory analysis with Menon<sup>28</sup> |
|  | 20 | A single Freedman ES derived organoid isolated from a full well | Har_F_SO |  |
| Subramanian *et al.*<sup>14</sup> | 7, 15, 29, 32 | Takasato iPS derived organoids with 3 pooled replicates per time using iPS cell line designated "ThF" | Sub_T_L1 | Clustering & DE genes, organoid trained random forest classifier, integration with Young<sup>29</sup>, Lindstom<sup>19</sup> and self-generated kidney tissue |
|  | 7, 15, 29 | Takasato iPS derived organoids with 3 pooled replicates per time using iPS cell line designated "AS" | Sub_T_L2 |  |
| Kumar *et al.*<sup>35</sup> | 25 | Modified Takasato iPS derived micro-organoid | Ku_TMO | Integration with organoid<sup>15,33</sup> with clustering & DE genes |
| Low *et al.*<sup>36</sup> | 10, 12, 14 | Modified Takasato ES derived organoids | Low_TMod | Clustering & DE genes |
| Tran *et al.*<sup>26</sup> | 16, 28 | Morizane ES derived organoids | Tran_M | Clustering & DE genes individually & after integrating with self-generated kidney tissue |
| Lawlor, Vanslambrouck, Higgins *et al.*<sup>37</sup> | 25 | Takasato iPS derived organoids generated by bioprinting. Organoids were compared with three different biophysical properties. | LVH_T | Clustering & DE genes compared to Hochane<sup>25</sup> trained machine learning model using scPred |
| Howden, Wilson *et al.*<sup>24</sup> | NA | Takasato iPS derived organoids dissociated and GATA3+EPCAM+ cells isolated. These cells cultured in ureteric epithelium promoting conditions. | HW_iUB | Seurat Label Transfer using reanalysed Holloway |
| Mae *et al.*<sup>38</sup> | NA | Induced Ureteric Bud cultures | Mae_iUB | Clustering & DE genes |

Here we have taken reference HFK datasets from three publications that span multiple ages and kidney regions, performed individual annotations of the cells present based on prior information, then used all confidently classified cells to train classification models using the *R* package *scPred*^22^, a generalizable method which has showed high accuracy in different experiments and datasets from multiple tissues, and considered a top performer in benchmarking studies^9^. The resulting model, referred to as *DevKidCC*, provides a robust and accurate classification of cells in novel single cell datasets generated from developing human kidney or stem cell-derived kidney organoids. *DevKidCC* defines a model of cellular identity organised in a hierarchical manner to represent the key developmental trajectories of lineages within the developing kidney. The classification method is complemented with custom visualization tools in the *DevKidCC* package. This classifier was then used to investigate published kidney organoid datasets to compare organoid patterning and gene expression profiles across these datasets. We present a variety of applications of *DevKidCC* to the reanalysis of existing data. This analysis revealed differences in nephron progenitor proportion and nephron patterning and maturation between kidney organoid protocols. We also apply *DevKidCC* to investigate approaches for directed differentiation to ureteric epithelium and dissect the effect of all-trans retinoic acid on nephron patterning and podocyte maturation. While *DevKidCC* is specifically trained on HFK for application to kidney organoid models, the framework presented here could be applied for any tissue system to generate a cell classification model.

## Results

### Generation of the model hierarchy for complete cell classification

We first build a comprehensive reference dataset on which to train the probabilistic classification models. We used single cell RNA-sequence datasets from three publications currently generated on HFK (Table 1). Samples range from 9 to 19 weeks’ gestation across which time the developing human kidney undergoes both growth and maturation, with week 16 being most frequently represented. Cells in all were originally annotated using clustering and cluster labelling using marker gene expression. One dataset was a recently published high quality HFK dataset^23^ (8,987 cells) that included both medulla and cortex regions and including a 96-day male and 108-day female sample. Of note, this dataset contained ureteric epithelium, which had not been thoroughly analysed to this point^24^. This data was combined with data from 17,759 HFKs cells ranging from week 11 to 18 of gestation^25^ to increase the developmental range of the training set. A further 8,317 cells from gestational week 17 which had been microdissected into cortex, inner and outer medullary zones^26^ were combined to complete the comprehensive reference single cell RNA-sequencing HFK dataset. Cells from all datasets were integrated using *Harmony*^27^ (Figure 1A) before performing a supervised clustering and annotation, using the original annotations of each dataset as a guide. This led to a reference dataset containing three ureteric epithelial subpopulations (Tip, OuterStalk, InnerStalk), four stromal subpopulations (Stromal Progenitor Cells (SPC), Cortex, Medullary, Mesangial), endothelium, the nephron progenitor cells (NPC) and the nephron including subpopulations of CellCycle (CC), EarlyNephron (EN), early distal and medial tubule (EDT_EMT), distal tubule (DT), Loop of Henle (LOH), early proximal tubule (EPT), proximal tubule (PT), parietal epithelial cells (PEC), early podocytes (EPod) and podocytes (Pod) (Supplementary 1). These populations have been further classified in the original publications, such as the DT being split into distal straight, distal convoluted and connecting segment or classifying populations in relation to morphological features, such renal vesicle, comma shaped body and S-shaped body segmentation^24–26^. While morphologically there is a consistency in segment identification, this is less clear in single cell data and has led to inconsistency in classification terminology. As such, here we have classified cell populations based on expression of known differentiation markers as cells take on a more distinct identity (Figure 1B).

**Figure 1:**
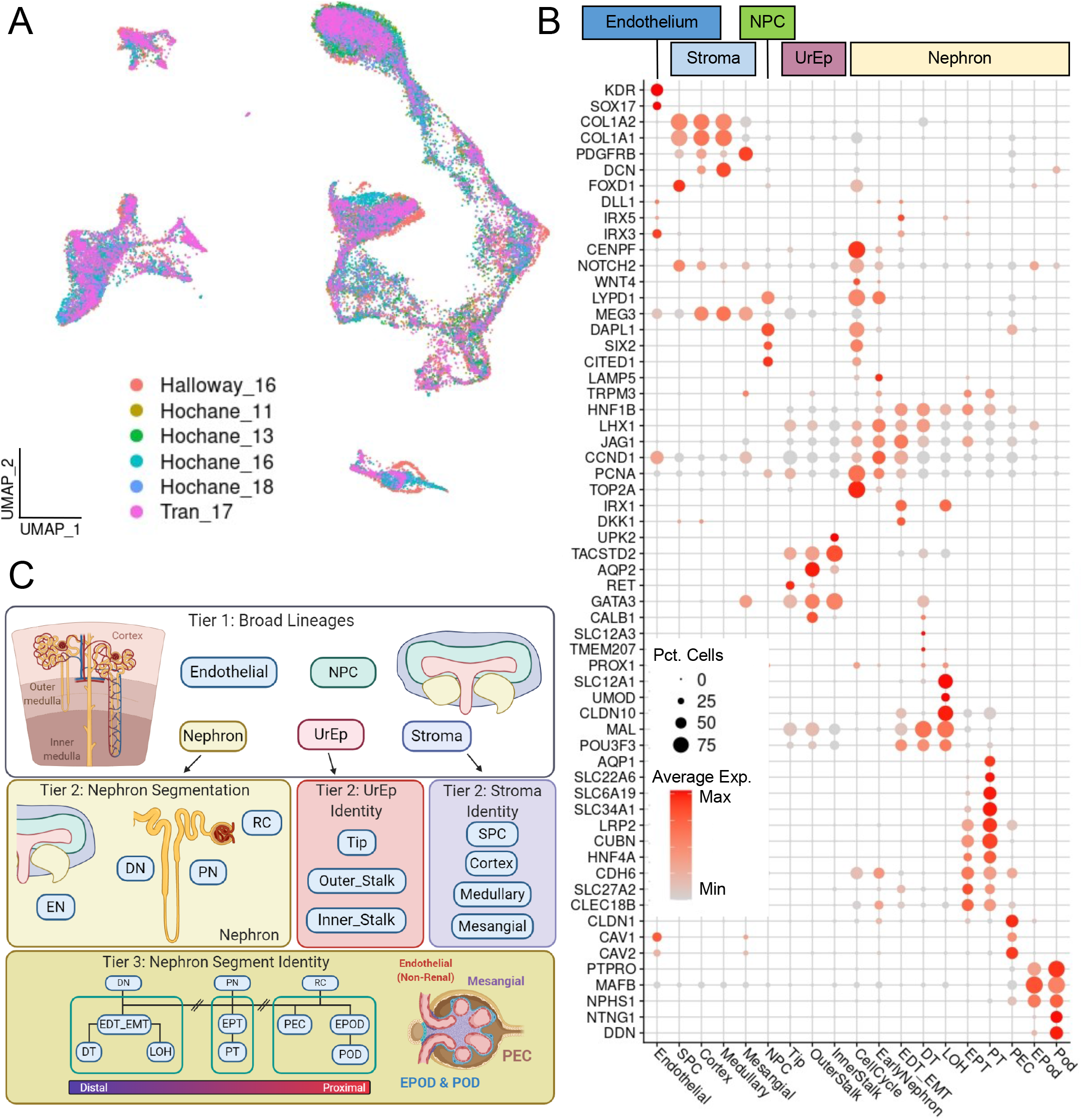
Generating a comprehensive reference to train the classification model hierarchy. A) The comprehensive human fetal kidney reference displayed in the first two UMAP dimensions grouped by their dataset of origin. The samples were integrated using *Harmony.* B) A DotPlot showing the expression of known marker and important genes present in each kidney segment. C) Graphical outline of the model hierarchy employed by *DevKidCC* to classify cells from single cell RNA sequencing kidney organoid datasets.

The complex and dynamic nature of the developing kidney, with multiple cell lineages and waves of nephrogenesis, means that cells of many stages of differentiation can be present at all developing timepoints within the same single cell data. This is one of the main challenges in classifying cells in the HFK single cell data, as the cells are in transitional flux. The multiple lineages within the kidney also make classifying cell types difficult, as the differences between lineages mask the subtle differences in gene expression between cell types within a lineage, such as those of the epithelial sub-types. To minimise the impact of this transcriptional variance on classification, we took a hierarchal approach by training three tiers of models (Figure 1C). The first tier classified cells based on their lineage; nephron progenitor cells (NPC), nephron, ureteric epithelium (UrEp), stroma and endothelial. The second tier for the UrEp lineage classified cells into the highly proliferative Tip cells, *AQP2*-expressing outer stalk and uroplakin-expressing inner stalk. The second tier for the stroma lineage classified cells into the *FOXD1*-expressing stromal progenitors (SPC), the cortical and medullary stroma clearly identifiable in the outer and inner zones^26^ (*DCN* low/high respectively) and the mesangial cells which express *GATA3*. The nephron segmentation required an extra tier due to the complexity of cell types present and their transcriptional similarity. Here, the second tier classified the early nephron (EN) that could not be clearly identified as polarised, the proximal (PN) and distal (DN) nephron epithelium and the renal corpuscle (RC) lineage. These were then further classified in third tier models (Figure 1C). The models were trained with the package *scPred* ^22^ using a support vector machine with a radial basis kernel and 100 principal components. The *scPred* package utilises a machine learning approach to train predictive models on a reference single cell dataset. This model can estimate the similarity of a cell within a query dataset to the identities classified within the model. This has been shown to be a robust method to classify cells of a novel dataset based on a known reference^9,37^. We created wrapper functions of all the models into a single use function (*DevKidCC*), which takes an input of a *Seurat* object. To determine cells in the first tier, we use a probability threshold of 0.7, while at all other tiers the threshold is removed. This enables all cells that are classified at the top tier to be given an assigned identity regardless of the highest level of similarity predicted by the lower tier models. Further investigation of the calculated similarity value can be interrogated as every cell has a record in the metadata of the scores from each classification. No pre-processing is required as data is normalised during the function call. The recommended pipeline is to read in raw counts data using the *Seurat* pipeline, filter out poor quality cells and then run *DevKidCC*. The classifications for each tier and the final identities can be accessed within the metadata slot for further investigation. The package contains custom in-build functions *ComparePlot*, *DotPlotCompare* and *SankeyPlot* to investigate the cell populations within the classified sample.

### *DevKidCC* classification rapidly and accurately reproduces published annotations

While this tool was designed to classify cells within kidney organoids, we first confirm the capacity to accurately classify developing kidney cell types by applying it to other HFK datasets. We applied *DevKidCC* to the dataset of Lindstrom^19^ of which the original cell classification identities were equivalent to those of the first classification tier within *DevKidCC* (Figure 2A). *DevKidCC* classified 90% of the 2945 cells that passed quality control, while the remaining cells expressed markers for immune cells (*HLA-DRA*, *CCL3*, *SRGN*) which are not represented in the model and so were not assigned an identity. 14 cells were classified as UrEp, positioned at the tips of one end of the nephron cluster, which *DevKidCC* further classified as DN epithelium. While these two cell populations arise from distinct precursors, they share a very similar transcriptional profile, making them very difficult to distinguish at single cell level^15–18,24^. The ability to identify and classify these two populations separately, even with a small contribution of one population within a dataset, demonstrates the power of *DevKidCC* as a classification tool, particularly in comparison to clustering algorithms. The expression of marker genes used by Lindstrom^19^ to annotate cell identities were shown as enriched in the same populations classified using *DevKidCC* (Figure 2B), affirming the accuracy and relevance of our classification tool.

**Figure 2:**
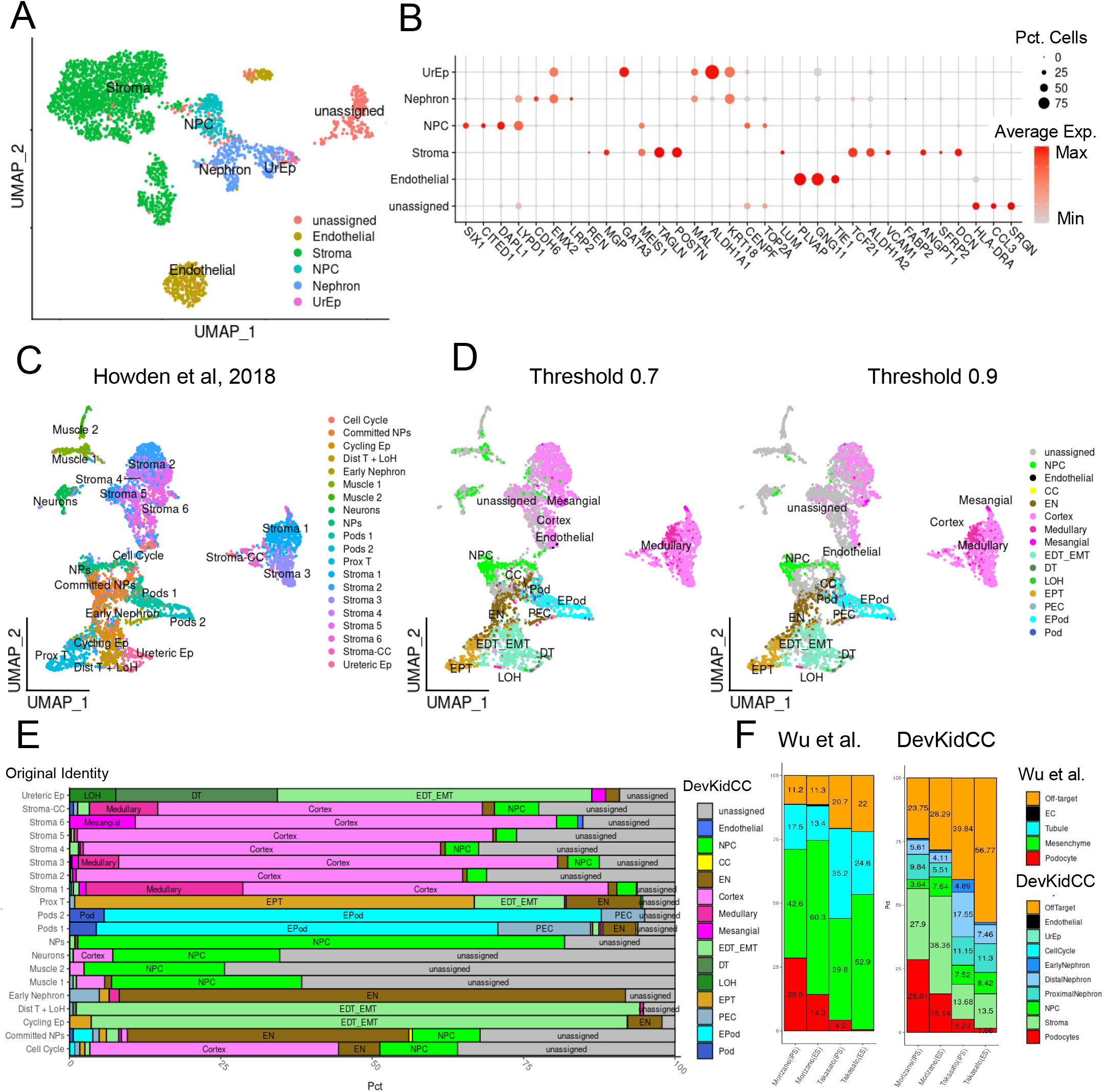
*DevKidCC* classification in human fetal kidney and organoid datasets. A) UMAP representation of the Lindstrom 2018 human fetal kidney dataset, grouped by the *DevKidCC* classification. B) A *DotPlot* showing the expression of key marker genes from the original Lindstrom 2018 analysis and their expression in the *DevKid* 13 *CC* tier 1 annotated cell types. C) UMAP representation of the original annotation from Howden kidney organoids. D) UMAP representation of cell classification using *DevKidCC* using thresholds of 0.7 and 0.9 similarity scores. E) A *ComparePlot* showing the reclassification of cells from the original Howden^13^ annotation using *DevKidCC*. Cell classification is well conserved when considering differences in nomenclature. F) Directly comparing the original annotation of four organoid samples from Wu^12^ to that of *DevKidCC* shows the congruence of classification with increased accuracy in determining kidney-like mesenchymal cells from non kidney-like cells.

The arbitrary nature of classifying cells using clustering algorithms is challenged when identifying cells transitioning between populations, often represented as the “borders” of clusters. The cluster-based classification of such cells will change with different approaches to analysis. The application of a cell-centered identification approach circumvents this challenge. To investigate this, *DevKidCC* classification of two published kidney organoid single cell datasets was compared to their original cluster-based annotations. Howden^13^ contained samples from two differentiation timepoints; intermediate (18 day) and late (25 day) stage organoids (Figure 2C) while Wu^12^ contained datasets from two distinct protocols for deriving kidney organoids, labelled as Takasato^30^ and Morizane^31^ after the original authors. *scPred*^22^ allows for the setting of a threshold of minimum similarity for a cell to be assigned a given identity. The distribution of the maximum scores for cells in the HFK and organoid datasets showed very similar patterns, however in the HFK there are more distinct peaks at the higher end of similarity (Supplementary Figure 2). In organoids we see a more gradual decrease in scores, meaning there is no set point at which the threshold should obviously be set (Supplementary Figure 2). Organoid datasets from Howden^13^ and Wu^12^ displayed a similar distribution of similarity scores for Stroma and NPC (Supplementary Figure 2). A sensitivity analysis was performed by comparing threshold points of 0.7 and 0.9 with the Howden^13^ dataset where we had access to the original annotation for each cell. When mapping the *DevKidCC* classification at both 0.7 and 0.9 thresholds onto the UMAP plot and comparing this to the original classification, *DevKidCC* accurately replicated the original annotation in both settings (Figure 2D). Only a small number of cells did not get classified at the top tier model, defining them as “unassigned” cell types. Such cell types may represent non-renal off target cell types not normally present in HFK or cells in which identity is not sufficiently strong to definitively classify. While all original clusters contained cells that were reclassified as unassigned, the largest contribution was from clusters previously annotated as neuron and muscle, illustrating the specificity with which the model classifies renal cell types (Figure 2E).

Both stroma and NPC are mesenchymal cell types. The mesenchymal cells present within kidney organoids have been difficult to accurately classify due to their gene expression profiles being different to those of characterized developing kidney stroma^15^. The previous analysis of the Howden^13^ dataset identified seven clusters as stromal (Figure 2C), of which almost all of those assigned an identity using *DevKidCC* remained classified as a stromal sub-type (Figure 2E). However, of the two largest stromal clusters initially identified (see Figure 2A^13^), the *TCF21^+^* cluster showed a higher number of classified stromal cells while the second was more ‘unassigned’. When looking at the similarity scores for stroma and NPC identity in tier 1, the *TCF21* expressing population showed a stromal similarity > 0.9 but very low NPC scores, while the other population had lower stromal scores, albeit still predominantly >0.7, and higher NPC scores, indicating these represent a less defined mesenchymal population (Figure 2D, Supplementary Figure 2). Within the nephron, cells previously identified as “Committed and Early Nephron” were reclassified by *DevKidCC* to comprise a smaller population of NPC together with a larger population of cells identified as Early Nephron (Figure 2E). To examine this further, we analysed organoids generated from either embryonic (ES) or induced pluripotent (iPS) stem cells using two different protocols^12^. Using *DevKidCC* we were able to rapidly reproduce the initial classification of these organoids, accounting for the differences in the nomenclature (Figure 2F). Using *DevKidCC* classification we identified cells which do not match the reference (termed “unassigned”) enabling further investigation. Here, *DevKidCC* could again distinguish kidney stroma from likely off target cell types like muscle and neural that may represent artefacts of *in vitro* culture^12,13^. Together this reanalysis demonstrates the accuracy with which *DevKidCC* can classify renal cell types within organoid datasets.

### *DevKidCC* provides a method for direct comparison between protocols

A major challenge for the field has been to compare between datasets generated from different labs, lines, batches or from different protocols due to differences in the analyses that were used. This is particularly pertinent given the use of several distinct protocols for generating kidney tissue from hPSCs (Takasato^30^, Morizane^31^ and Freedman^39^, see Table 1). Direct comparisons between studies and protocols requires an integration of all existing samples to allow re-clustering and differential gene expression analysis on the combined dataset. This is challenging due to the noise between samples, the majority of which relates to technical or batch effects^33^ which can confound biological variations of interest during data integration^40^. To avoid these challenges, *DevKidCC* was used to directly identify all cell types present within multiple datasets enabling direct comparisons without the need for integration. As *DevKidCC* will compare all cells to the same comprehensive reference, the biological information for each sample can be directly compared without prior dimensional reduction and clustering. To demonstrate this, we applied *DevKidCC* to all available single cell kidney organoid datasets (summarised in Table 1) irrespective of the cell line, organoid age, differentiation protocol or laboratory. The resulting comprehensive analysis (Figure 3) allows a direct comparison of cell proportions across all samples at each tier of classification, grouped into the three main differentiation protocols represented in the literature (Figure 3). What is immediately evident is both the variation in the proportions of “unassigned” cells across all datasets and the lack of nephron maturation even in the oldest organoids regardless of protocol. The maturation of nephron cell types was limited in all protocols and samples, although the Morizane^31^ protocol produced organoids with the highest number of cells reflective of a more mature podocyte stage. While there are a small number of mature podocytes, there are almost no mature proximal tubule cells generated with any organoid protocol, but rather being classified as less mature EPT. These have expression of proximal markers such as *CUBN*, *LRP2* and *HNF4A* but lack the specific solute channels such as *SLC47A1*, *SLC22A2* and *SLC22A8* (Supplementary Figure 3). In clustering-based analyses, these cell populations are often split into two or more groups which are interpreted to have varying degrees of maturation, whereas the *DevKidCC* classification indicates that these are mostly immature.

**Figure 3:**
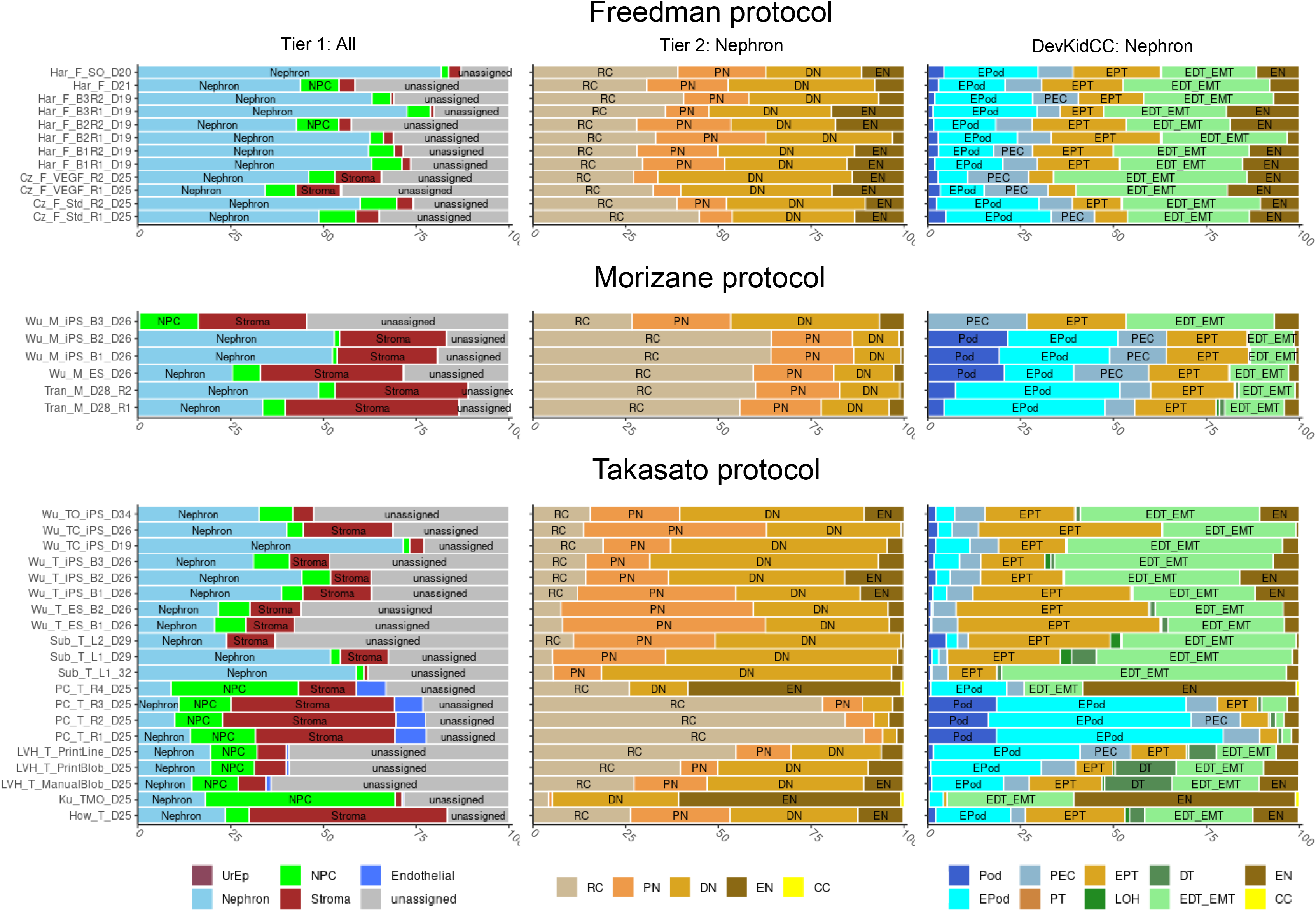
Direct comparison of organoids generated from different protocols. All the mature age organoids were classified using DevKidCC. The proportions of identities are classified, the first column is all cells classified at the top tier, the second column is the nephron cells classed at the sec by the ond tier, and the last column is the nephron cells classified at the third tier. Samples are grouped in rows protocol used to derive the organoids.

*DevKidCC* analysis revealed differences in cell proportion and nephron patterning between organoids generated with different protocols. Organoids generated using the Freedman^39^ protocol show a small stromal population in comparison to other protocols. The Morizane^31^ organoids show little early nephron cell identity while the Freedman^39^ organoids tend to have more early-stage nephron cells. In the Morizane^31^ organoids we identify limited distal tubule regions, having less than 25% of the nephrons classified as distal whereas in the Takasato^30^ and Freedman^39^ protocols is more evenly segmented across nephron components. The Takasato^30^ protocol generates the most distal tubule, including some cells classed as a more mature DT segment as well as a Loop of Henle population (Figure 3). The DT expressed *GATA3* and *TMEM52B* but lacked the distal convoluted tubule (DCT)-specific marker *SLC12A3*. However, in some cases the connecting segment (CS)-specific marker *CALB1* is expressed. This would indicate that the connecting segment, which represents the most distal region of the nephron and which invades and fuses into the ureteric tip to form a contiguous tube, is being generated in some organoids. This is promising as it would indicate that there is the potential to promote fusion of these nephrons to any separately induced collecting duct structure, potentially through engineering methods. In summary, while nephrons are forming and showing evidence of patterning and identifiable segmentation in all protocols, one should keep in mind their relative proximo-distal patterning and evident immaturity prior to their application in disease modelling and drug screen studies.

### Identifying nephron progenitor cell variation between protocols using ComponentPlot and *DotPlotCompare*

To further investigate relative gene expression between datasets, we extracted gene expression profiles and proportions of cells in each classified population, in all available organoid datasets (see Table 1) and the comprehensive reference. A modified version of the *DotPlot* function from the *Seurat*^7,8^ package was included to compare gene expression between datasets. The direct comparison between kidney organoids (Figure 3) revealed substantial variation in the proportion of NPC, which we further investigated applying the modified function named *DotPlotCompare* to visualization relative gene expression in NPCs across all protocols.

The nephron develops from NPCs which are a heterogeneous population of mesenchyme that undergo a mesenchyme to epithelial transition (MET) in response to signals from the ureteric epithelium, giving rise to the entire nephron epithelium^41,42^. *In vivo* analysis has shown markers like *SIX1*, *SIX2*, *CITED1*, *DAPL1* and *LYPD1* are expressed in this population and can be used to reliably identify these cells from the surrounding stromal mesenchyme *in situ*^17,19^. These markers have also been used to identify the NPC populations of cells in both HFK and organoids in single cell datasets. NPCs express a posterior HOX code, particularly the HOX10 and HOX11 paralogues^43,44^. Visualising the NPC populations from within the reference HFK dataset using *DevKidCC*, we can see that 44.9% of cells express *SIX2*, 56.3% express *SIX1*, 53.3% *CITED1* while over 70% express *DAPL1* and *LYPD1* (Figure 4A). The posterior HOX genes are also expressed, with *HOXA10* most abundant and *HOXC10*, *HOXD10*, *HOXA11* and *HOXD11* at lower levels and in less cells (Figure 4A). The surprising heterogeneity of gene expression within this population could be explained by technical challenges, including data sparseness, dropout levels and capture bias. It may also be explained by transcriptional bursting^45^, where genes are not constantly being transcribed and so the sample harvesting may occur during a transcriptional lull. However, this does provide a true reference for comparison to the expression profiles expected within these cell populations in organoids.

**Figure 4:**
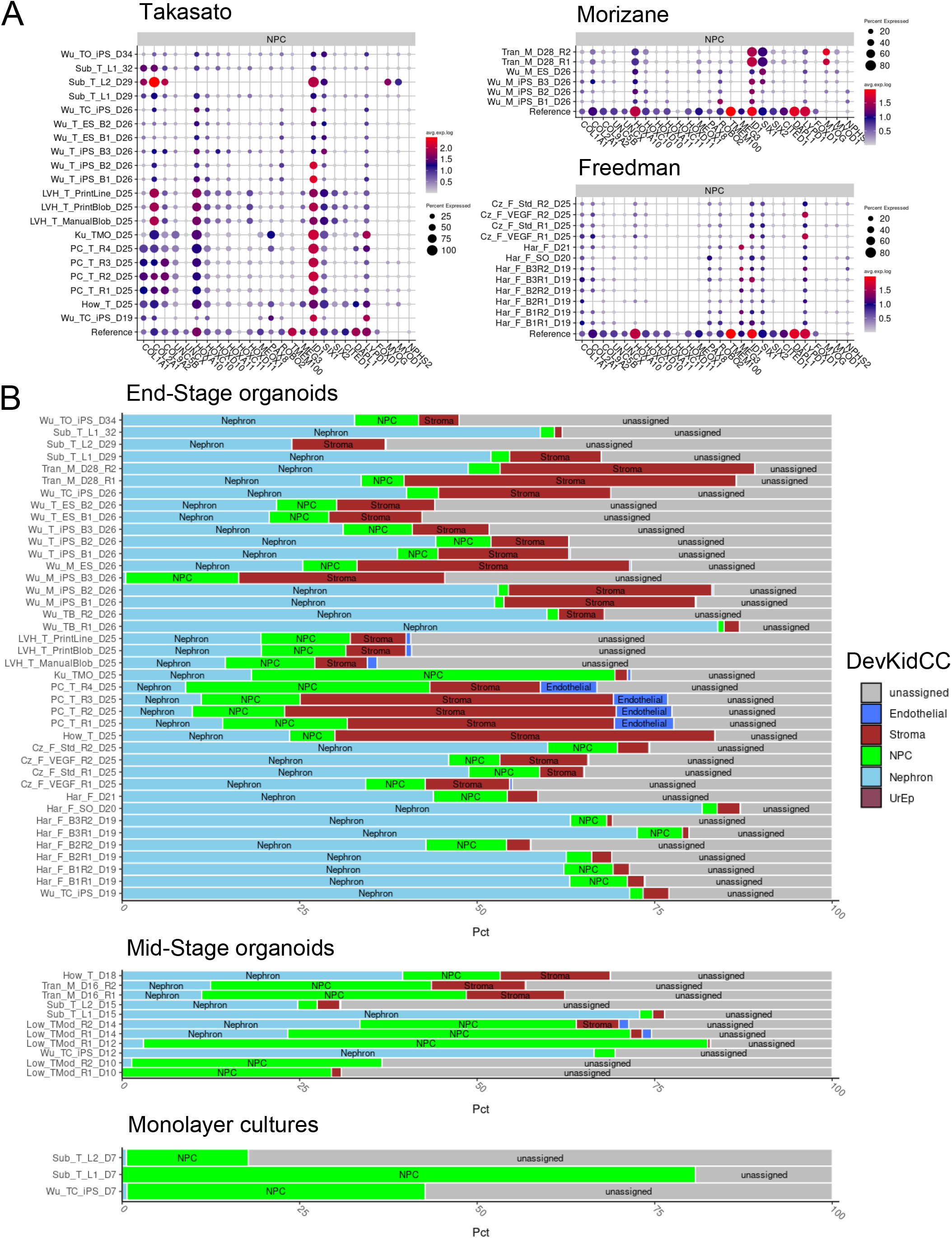
NPCs deplete as organoids age and vary in transcriptional similarity to HFK NPCs. A) Expression of NPC marker genes in the NPC cluster from each sample of each publication compared to the reference dataset. Takasato derived organoids show more congruence with the reference profile than Morizane or Freedman organoids. B) Proportion of tier one classification in i) organoids 17 days or more, ii) organoids 16 days or less, iii) monolayer differentiations showing the variation of NPC contribution across ages and datasets.

When we compare organoid NPCs to the HFK reference, we again note variance between publications and protocols. While NPCs constitute 5-10% of the total cells for the Freedman protocol (Figure 3, 4B), these populations have almost no expression of *SIX2*, *CITED1* and *DAPL1*. Similarly, they do not express the posterior HOX code, and only express low levels of *LYPD1* and *SIX2* (Figure 4A). Organoids generated from the Morizane protocol have a more similar profile to the reference NPCs, including some posterior HOX gene expression but little *SIX1* and *LYPD1* expression and almost no *SIX2*, *CITED1* and *DAPL1* present. Takasato-derived organoid NPCs have the most similar profiles to the reference, with NPCs in some samples coexpressing *SIX1*, *SIX2*, *CITED1*, *DAPL1* and *LYPD1*. There is some variance between publications generating organoids from the same protocol (Figure 4B), concurring with earlier studies showing that batch differences are a notable source of variation^12,33^. In the “unassigned” populations generated in organoids, expression of the muscle markers, including *MYOG* and *MYOD1* was sometimes evident. A subset of individual cells within such a published ‘muscle’ cluster^13^ were re-classified by *DevKidCC* as NPC but do show expression of these muscle genes (Figure 4A). Indeed, muscle gene expression is detectable in kidney organoid clusters previously labelled as NPC from multiple protocols and publications^12–15,32^. However, there is no evidence the expression of these genes in the HFK reference, suggesting that their consistent expression in organoid populations is an artifact of the *in vitro* culture conditions. This demonstrates how using *DevKidCC* to classify and directly compare all published organoids datasets can improve our understanding of NPC population generated across multiple kidney organoid protocols. We have identified an *in vitro* culture artefact muscle gene signature within the NPC population present across multiple protocols, giving a target to modulate for improving NPC identity within organoids.

### *DevKidCC* classification highlights distinct expression profiles of organoid podocytes compared to human fetal kidney

Estimating the maturity of cells within a single cell dataset is commonly performed by combining an analysis of cell specific maturation markers within clusters and placing cells or clusters along pseudotime trajectories. However without a time-stamped reference to align the transcriptional profiles these results can be open to interpretation. *DevKidCC* classifies cells based on a reference dataset with a range of maturation states, enabling us to directly compare maturation levels across samples.

The glomerular epithelial cells or podocytes are a non-dividing architecturally constrained cell type surrounding the fenestrated capillaries within the glomerulus of each nephron. Podocytes arise from the proximal nephron with trajectory analysis suggesting a distinct transition from the NPCs to that of the remaining nephron epithelium^12,25,28^. Forming a component cell type of the renal corpuscle / glomerulus, the podocytes are anatomically surrounded by a Bowman’s capsule comprised of parietal epithelial cells (PECs) which show transcriptional overlap with both podocytes and proximal tubule^46^. Hochane^25^ defined a pattern of differential expression across podocytes during maturation. Here, *OLFM3* was expressed in the proximal end of the early nephron (S-Shaped body stage) preceding podocyte patterning with expression of this gene decreasing during podocyte maturation and upregulation of *NPHS1* and *NPHS2*. While expressing markers of podocyte at a lower level, including *MAFB*, *TCF21*, *NPHS1* and *NPHS2*, PECs showed specific expression of *CLDN1* and enriched expression of *PAX8*^24,25,47^. *DevKidCC* analysis of organoid protocols classified most renal corpuscle components as immature podocytes (EPod), with most protocols containing cells classified as PEC and EPod. Organoid EPod and Pod populations had varying levels of *CLDN1*, while *OLFM3* and *PAX8* were co-expressed with more mature podocyte markers like *NPHS1* and *NPHS2* in the PEC and Pod populations. (Figure 5). This may indicate that *in vitro* podocyte differentiation does not progress in the same manner as *in vivo* or that these cells are undergoing maturation. The key collagen genes expressed by the podocytes to form a mature glomerular basement membrane are *COL4A3*, *COL4A4* and *COL4A5*^48^. Organoid podocytes again show low expression of these genes compared to podocytes in the HFK reference data. The exception to this observation was seen in organoids subjected to a longer period of time in culture^14,33^ suggesting a capacity to mature with time. A critical switch in podocyte maturation is suppression of proliferation, with this post-mitotic state maintained via the expression of key cell cycle regulators including *CDKN1A* (p21) and *CDKN1C* (p57). This is seen in the reference with an increase in expression in the EPod and Pod populations, paired with a decrease in mitotic markers such as *TOP2A,* however in the organoid podocytes there is little decrease in mitosis markers, but expression of *CDKN1A* and *CDKN1C* do increase (Supplementary Figure 4). As such, *DevKidCC* can also be employed as a tool to gain biologically relevant insights into kidney organoids generated from different protocols and users. This is promising for the application of such a tool to compare between wildtype and mutant organoid datasets.

**Figure 5:**
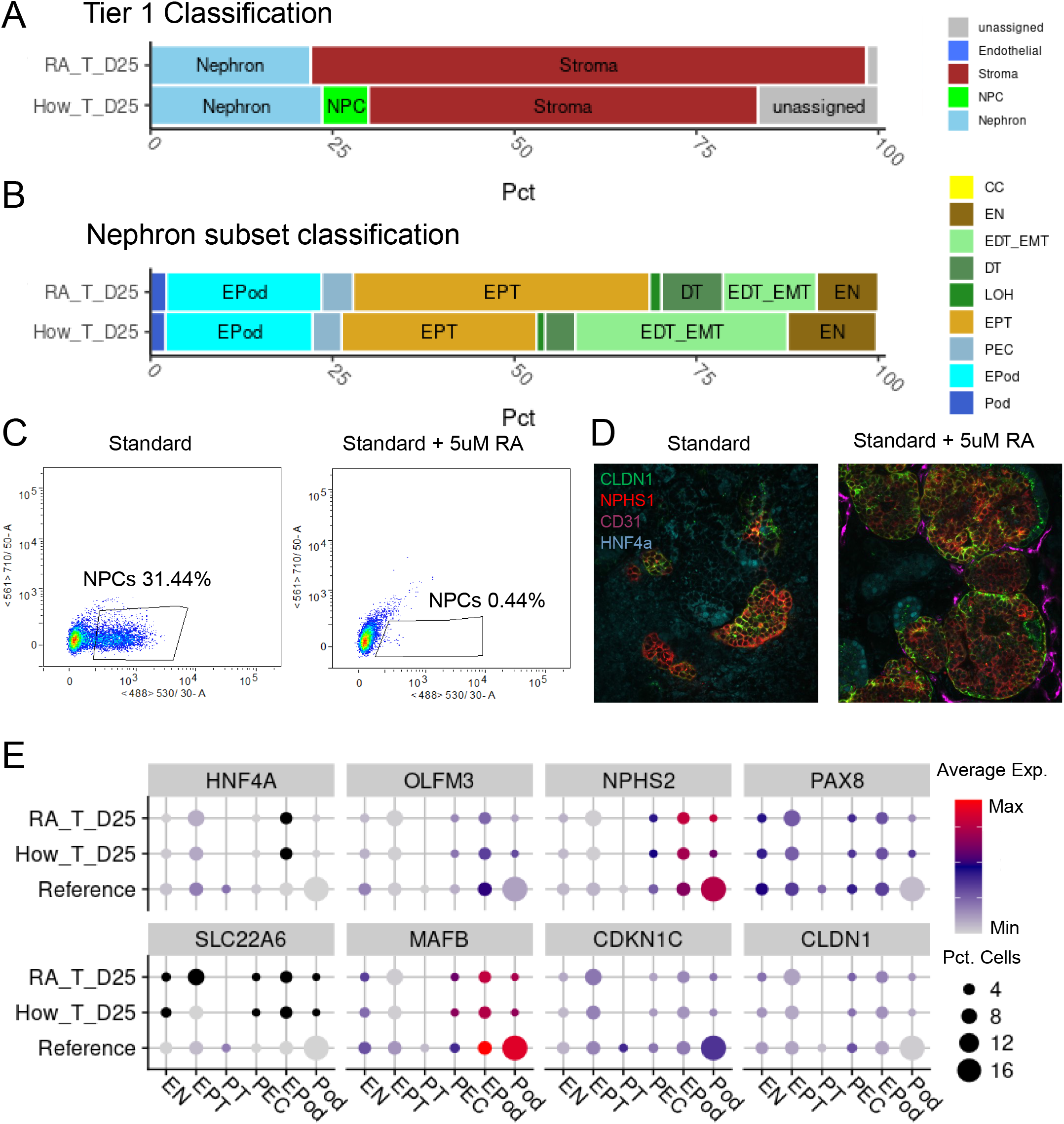
Addition of Retinoic Acid during organoid development depletes NPCs and promotes glomerular maturation. A) The proportions of cells classified between day 25 organoids of standard protocol or with Retinoic Acid (RA) added at D7+5 (D12). B) The proportion of nephron cells classified into their subpopulations shows an expansion of the proximal tubule at the expense of the distal tubule and early nephron when adding RA. C) FLOW plots showing the expression of SIX2^EGFP^ cells from D7+11 (D18) organoids with and without RA addition at D7+5 (D12) of protocol. D) The expression and localisation of CLDN1 can clearly be seen to be improved in RA organoids by immunofluorescence. E) Comparative gene expression between the reference, standard organoid and RA organoid for informative genes. *HNF4A* and *SLC22A6* are expressed in immature and mature proximal tubule respectively. *OLFM3*, *MAFB* and *NPHS2* are expressed in precursor, immature/mature and mature podocytes respectively. *CDKN1C* is a post-mitotic marker. *PAX8* is expressed in the nephron epithelium but not mature podocytes. *CLDN1* is expressed in the parietal epithelial cells.

### Application of *DevKidCC* to investigate the impact of retinoic acid on kidney organoid maturation

Accurately identifying the cell types present within an organoid is crucial for the analysis of disease states or the optimization of the differentiation protocols. To evaluate the application of *DevKidCC* in analyzing functional differences between methods, we analysed unpublished data in which kidney organoids from the same starting cell line generated in the same batch were treated with 5μM retinoic acid after removal of all other growth factors at day 12 (7+5) of the protocol to promote maturation. Mammalian nephrogenesis *in vivo* occurs in waves with new nephrons constantly forming up to 36 weeks gestation^49,50^ in humans and into the first week of life in mice^51^. This is facilitated by the presence of a peripheral nephrogenic niche within which the NPC balance self-renewal versus nephron commitment. Once differentiated, NPCs exist throughout the duration of organoid culture and deplete with time, although a population does remain in mature organiods able to undergo nephrogenesis when induced with a canonical Wnt agonist^13^ (Figure 4A). Retinoic acid signaling plays many roles in kidney development depending on spatiotemporal expression^52–54^, and is also known to promote the differentiation of progenitor cell populations^55^. We investigated adding all-trans retinoic acid (RA) to organoids at multiple time points to see what effect this would have on organoids. The addition of 1-5 μM RA before day five of 3D organoid culture, substantially impaired nephron formation, whereas addition at day five onwards led to organoids with fully segmented nephrons similar to organoids without RA (data not shown). The *DevKidCC* classification identified an increase in the percentage of classified stromal cells, seemingly at the expense of the ‘unassigned’ population. In contrast to control organoids at day 25, the addition of RA resulted in a complete depletion of NPC cells (Figure 5A). While the percentage of nephron cells did not change, there was a shift towards more proximal tubule (EPT) than early distal tubule (EDT_EMT) (Figure 5B). These comparisons show evidence that RA caused the depletion of NPCs and proximalisation of nephrons within forming organoids. NPC depletion can be seen 6 days after addition of RA, when organoids generated using a SIX2^EGFP^ reporter line^13,56^ were analysed by flow cytometry. The control organoids had 31.44% EGFP+ cells while the organoids with RA had less than 0.5% (Figure 5C). This confirms that RA acts directly or indirectly on the NPC population, forcing them to either undergo commitment to form nephrons or differentiate away from NPC identity down a stromal pathway.

To investigate the maturation of the nephrons we visualized maturation markers for all segments using the *DotPlotCompare* function within the package. Only the podocytes showed evidence of maturation, with an increase in the expression of genes such as *WT1, MAFB, TCF21* and *NPHS2* with RA addition while also showing a decrease in *OLFM3* expression, a marker of the immature podocytes^25^. Interestingly, the PEC marker *CLDN1* remained expressed in the podocytes, although immunofluorescence showed more specific localization to the epithelial cells surrounding the podocytes, which is the normal location of PECs (Figure 5D, 5E). These results may indicate that the podocytes and potential PEC cells increase in maturity when RA is added. The expression of both PEC and podocyte markers in cells assigned to all three renal corpuscle identities is consistent with the previous analysis of these populations and may indicate that while maturation is occurring, the delineation of specific gene signatures within these cells is not.

### Analysis of existing protocols for the development of ureteric epithelium

*DevKidCC* is able to predict stromal, endothelial, ureteric and nephron cell identity based upon the reference data from HFK. Our analysis of existing standard organoid protocols confirms the absence of populations classified as ureteric epithelium. The ureteric epithelium in the mammalian kidney arises as a side branch of the mesonephric duct that grows into the presumptive kidney mesenchyme. Hence it has been suggested that it is not possible to generate ureteric epithelium using the same differentiation protocol able to generate the nephron lineages^57^. To date, a number of protocols have been published that report the generation of ureteric epithelium^24,38,57,58^ with all of all these methods involving the isolation of cellular fractions that are then cultured separately to form ureteric epithelium. Single cell analyses have recently revealed the significant transcriptional congruence between the distal nephron and the ureteric epithelium in both human and mouse^12,16^. It has also been established that distal nephron from standard organoids remains plastic and can be induced to adopt a ureteric epithelial fate^18^. To investigate how accurately *DevKidCC* can identify this kidney cell identity, we applied this analysis to a single cell dataset available from a specific UE protocol^38^ and the single cell dataset recently generated from UE that had been derived from DN^24^. *DevKidCC* classified 20% and 28% of cells as UE respectively (Figure 6A), with most of these classified as Tip (Figure 6B). The DN-derived sample^24^ contains a population of cells classed as nephron while the UE sample directly differentiated from hPSC^38^ contains cells classed as NPC (Figure 6A, 6B). The different cell types present between these two samples may be explained by the different protocols used to generate UE and kidney developmental biology. Cultures differentiated towards an anteriorised intermediate mesoderm population directly from hPSCs are likely to generate a proportion of NPC-like cells as a *bona fide* posterior intermediate mesoderm of a more anterior nephrogenic cord. In contrast, the DN-derived cultures contain nephron-like cells that have not become UE. In both samples the overwhelming majority of cells were “unassigned” (Figure 6A). However, when visualizing the distribution of scores there is an even distribution of UrEp scores in both samples between 0.1 and 1 (Figure 6C) with the majority of cells being most similar to the ureteric population over any other lineage. This indicates a spectrum of similarity to the true ureteric epithelium. The implications of this when attempting to classify these cells are that a clustering analysis will break them up without appreciating the overall transcriptional similarity while *DevKidCC* will classify each cell based on its own merit, giving a more accurate overall picture of cell identity compared to the true HFK profile.

**Figure 6:**
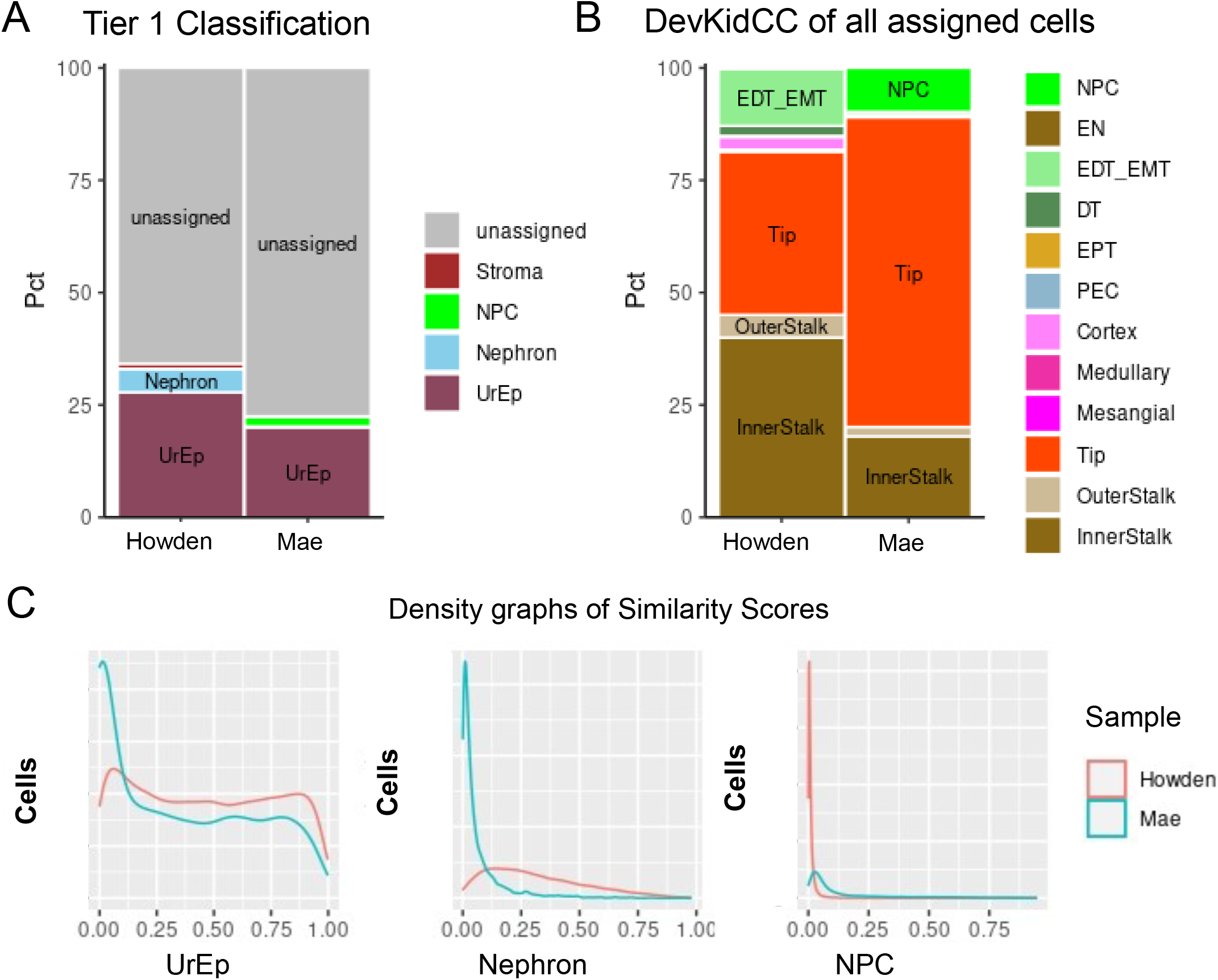
Classification of Ureteric cell types in targeted cultures. A) The *DevKidCC* classification for the Howden^24^ and Mae^38^ datasets show populations of ureteric cells (UrEp) classified. B) The complete classification of all cells not classed as “unassigned” shows interesting differences between the cell types present between the two datasets. Howden^24^ has a mix roughly 50:50 split of Tip to Stalk cells of the ureteric cells classified, and also distal nephron cell types. Mae^38^ has a higher proportion of Tip compared to stalk cells, and the nephron cell types are NPC not nephron epithelium. C) Density plots showing the spread of similarity scores for the UrEp, Nephron, NPC at the highest tier and the Tip and InnerStalk of the UrEp subsets.

## Discussion

The question of cell identity is one that is difficult to answer. Histologically we can try to define a cell type based on its morphology, gene expression or protein expression, the latter typically being read by immunohistochemistry and immunofluorescence assays. In many cellular states, particularly those present during organogenesis, evaluation of cellular identity by functional assays is challenging and marker expression is rarely unique. This challenge is significant when evaluating cell identity using single cell RNA sequencing data. Such data is sparse, providing an incomplete snapshot rather than a comprehensive picture. As capture technology and bioinformatics tools have improved, increased levels of information can be extracted from this data, providing an overall synergy of expression profile for groups of cells within a sample. This can be combined with the pseudotime trajectory or even molecular lineage tagging to relate cells within a sample by history, assisting in likely classification of cell type. Such inferences are much more difficult in a synthetic *in vitro* system such as hPSC-derived organoids. Such protocols direct cells to undergo a series of changes that attempt to replicate the *in vivo* process. However, in reality, hPSC-derived lineages often do not completely recapitulate their *in vivo* counterparts, at least at the level of the transcriptome. We can often identify a gene, or a number of genes, that are expressed in a cell that give us some information of what it can be classified as, but in many cases there is ambiguity. This is compounded by our knowledge that hPSC-derived organoid models replicate early developmental cell states that are frequently in flux, not present in adult tissue and are less well defined.

The classification of cells within all single cell data has been inconsistent as clustering and classification decisions vary between individual researchers and the limitations within each dataset. *DevKidCC* represents a method of specifically classifying individual cellular identity within hPSC-derived kidney organoids based upon models trained on a comprehensive reference dataset. Our tool facilitates direct comparisons between kidney organoid datasets by classifying cells based on the reference data. While the base package, *scPred*^22^, includes a way to integrate the data within the models using *Harmony*^27^, this can introduce false correlations between similar cell populations such as the mesenchymal cells that have intermediate to high scores for both stroma and NPC. Hence, *DevKidCC* provides an option to run the harmonization step, but this is not required or recommended for kidney organoid datasets. The classification for all datasets has been integrated into functions allowing for plotting any novel dataset in direct comparison using the classification from *DevKidCC*. Gene expression can be visualised using the *DotPlotCompare* function, while sample annotation can be visualised using *ComponentPlot* or *SankeyPlot.* These tools included in *DevKidCC* provide a classification and visualization toolset to investigate cell identity and gene expression within novel and existing kidney organoids.

*DevKidCC* was developed so that it could be applied to novel datasets facilitating direct comparisons to those previously generated. This will make comparative studies much easier, facilitating the analysis of genetic variants, disease states or methodological variation in new protocols. While this system has developed a model with three tiers of subclassification, the complexity of the human nephron, even in the fetal kidney, is such that there is scope to interrogate individual cellular identity even further within this and other subcomponents. As these models were trained using developing HFK, the ability of the tool to accurately classify cell identity during earlier stages of mesoderm patterning or mature kidney is limited. The adult kidney shows significant specification of functional cell types within all segments of the final nephron, many of which have distinct functional roles in renal filtration and fluid homeostasis but are not present in the fetal organ. Indeed, the ratio of epithelium to stroma is dramatically shifted in the adult. While the fetal kidney begins to form some more mature cellular states, such as the intercalated and principal cells of the distal nephron / collecting duct, it is likely that a distinct cellular identity tool will be required for the accurate identification of cellular identity in postnatal kidney tissue. Conversely, the use of HFK from Trimester 1 and 2 as the reference dataset limits the ability to identify earlier stages of morphogenesis. This may explain the large percentage of unassigned cell calls in datasets in early stages of kidney organoid differentiation protocols (Figure 3, Figure 4A). However, *DevKidCC* applied to early-stage differentiations (day 7, intermediate mesoderm) split cell identity between NPC and unassigned, suggesting that the tool is able to identify those cells beginning to commit to the mesenchymal precursors of the kidney. Indeed, in a dataset that includes day 7, 15 and 29 organoids between two cell lines^14^, there is a direct relationship between the proportion of cells classified as NPC at day 7 to the proportion of nephron cells at day 15 and 29 (Figure 4A). We conclude that at this early stage the cells identified as NPC at this early stage could be the percentage of the differentiation correctly patterned to intermediate mesoderm and are still the cells that will go on to form the nephron population.

## Conclusions

DevKidCC provides a robust, reproducible and computationally efficient tool for the classification of kidney single cell data, in both human and organoid-derived tissue. Using DevKidCC we can now directly compare between kidney samples regardless of batch and have done so for all available published datasets. This important advance has provided insights into differences in organoids derived using different protocols and allows for any novel dataset to be directly compared to all previous datasets. The included custom functions simplify visualisation of cell identity proportion and gene expression within samples and between multiple samples. Any novel dataset can be classified using the framework provided in this package, allowing for direct comparison to all previous datasets, all of which are included within the package. For visualisation of gene expression profiles and organoid cell identities, the gene expression profiles of all datasets have been built into an *R* Shiny app available at https://sbwilson91.shinyapps.io/devkidcc_interactive/^59^ that does not require the use of *R* directly, allowing for easy access to this information. Finally, while this package has been built using HFK data to classify kidney cells, the framework can be transferred to any tissue type where adequate single cell data is available.

## Methods

### *DevKidCC* algorithm

*DevKidCC* (Developing Kidney Cell Classifier) is a function written in *R* designed to provide an accurate, robust and reproducible method to classify single cell RNA-sequencing datasets containing human developing kidney-like cells. The algorithm has two steps: data pre-processing and cell classification. Below we describe the development and utilisation of these steps.

### Data pre-processing

The required input is a scRNA-seq dataset as a *Seurat*^7,8^ object. This object is first normalised by dividing the total expression of each gene by the total gene expression per cell then multiplied by a scale factor of 10,000 and natural log-transformed with pseudocount of 1.

### Cell classification

We generated a comprehensive developing kidney reference single cell dataset by harmonising the raw data from multiple high quality human fetal kidney datasets. The annotation of the reference included three tiers with increasing specificity, with a clear hierarchical structure between the tiers. This dataset was then used to train machine learning models using the *R* package *scPred*^22^. One model was trained for each node of identities within the classification hierarchy.

Utilising *scPred*^22^ the models were trained using the same parameters, with the relevant cells inputted for each. The feature space used was the top 100 principal components. The models were trained using a support vector machine with a radial kernel. The models are stored as a *scPred*^22^object and can be used to classify cells within a *Seurat*^7,8^ object using the *scPred*^22^ package. For classification, these models will calculate the similarity of a cell to each of the trained identities within that model, giving a probability score between 0 and 1 for each identity. It will then assign an identity of the highest similarity score above the set threshold, or call the cell unassigned if no identity scores above the threshold.

Cells are classified using these models, organised in a biologically relevant hierarchy so as to optimally and accurately identify the cellular identity of all analysed cells. All cells are first classified using the first-tier model, contains generalised lineage identities of stroma, nephron progenitors, nephron, ureteric epithelium and endothelium. After similarity calculation using the first-tier model, cells that do not pass the threshold are classified as unassigned. The threshold is set to 0.7 by default but can be adjusted by the user, which can be useful if the user wants to classify cells with at decreasing levels of similarity. Cells assigned to stroma, nephron and ureteric epithelium are passed into a second tier of classification specific to these identities. It is important to note that at the second and third classification tiers, there is no thresholding, i.e., all cells are assigned an identity with no cells classed as unassigned. The second-tier ureteric epithelium model is trained on the tip, cortical, outer and inner medullary cell identities. The second-tier stroma model is trained on the stromal progenitors, cortex, medullary and mesangial cell identities. The second-tier nephron model is trained on the early nephron, distal nephron, proximal nephron, renal corpuscle and nephron cell cycle population. The distal nephron, proximal nephron and renal corpuscle are then further classified into more specific identities in a third tier of models. The third-tier distal nephron model is trained on early distal/medial cells, distal tubule and loop of Henle cells. The third-tier proximal nephron model is trained on early proximal tubule and proximal tubule cells. The third-tier renal corpuscle model is trained on parietal epithelial cells, early podocytes and podocytes. Each stage of the classification step is recorded as a metadata column, as is the final classification for each cell. All the similarity scores and tier classifications are readily accessible within the *Seurat*^7,8^ object for further analysis.

### Comprehensive reference generation

Raw data was downloaded from GEO database from repositories GSE114530^60^ and GSE124472^61^, or provided to us directly by the authors, since made available at EMBL-EBI ArrayExpress under accession number E-MTAB-9083^62^. The data as *CellRanger* output was read into *R* and processed using *Seurat*^7,8^ (v3.2.2), using *SCTransform*^63^ for pre-processing. Clustering and manual annotation was performed on each dataset individually, referring back to the original papers and using established markers enriched in clusters to classify each cluster. Once annotated, datasets were integrated using *Harmony*^27^ with 100 PCAs and 10000 variable features.

### Organoid gene expression database

A reference database of all available kidney organoid datasets (Table 1) was generated by running *DevKidCC*, extracting summaries of the gene expression information at each classification tier, and combining these into a database. This database can be used to directly compare gene expression between existing datasets, also novel datasets classified using *DevKidCC*. The link to download this database is available at the package Github repository^64^.

### Downstream visualisation functions

To facilitate data visualisation and analysis of *DevKidCC* classified datasets, three customised functions were included in the package. *DotPlotCompare* is a modified version of the *DotPlot* function from the Seurat package. A gene expression profile of the reference is present within the function and can be used for direct comparisons to an existing or novel dataset. There is an option to visualise the organoid database within this function as well, the downloading instructions for this are available at the package Github repository^64^. The proportions of cells classified using *DevKidCC* can be visualised as a bar chart using the *ComparePlot* function. This can also take as input a gene and show the expression of that gene in each segment. The *SankeyPlot* function utilises the *networkD3* package to generate an interactive Sankey chart showing the flow of cell classification.

### DevKidCC Kidney Organoid Gene Explorer shiny app

To make visualisation of the organoid database possible outside of using *R,* a shiny app was developed^59^. This allows for an interactive way to visualise and analyse gene expression within published organoid datasets.

### iPSC-derived organoid differentiation

The day prior to differentiation, cells were dissociated with TrypLE (Thermo Fisher Scientific), counted using a hemocytometer, and seeded onto Laminin 521-coated 6-well plates at a density of 50 × 10^3^ cells per well in Essential 8 () medium. Intermediate mesoderm induction was performed by culturing iPSCs in TeSR-E6 medium (Stem Cell Technologies) containing 4-8 μM CHIR99021 (R&D Systems) for 4 days. On day 4, cells were switched to TeSR-E6 medium supplemented with 200ng/ml FGF9 (R&D Systems) and 1 μg/ml Heparin (Sigma-Aldrich). On day 7, cells were dissociated with TrypLE, diluted fivefold with TeSR-E6 medium, transferred to a 15-ml conical tube, and centrifuged for 5 min at 300 x g to pellet cells. The supernatant was discarded, and cells were resuspended in residual medium and transferred directly into a syringe for bioprinting. Syringes containing the cell paste were loaded onto a NovoGen MMX Bioprinter, primed to ensure cell material was flowing, with 100,000 cells deposited per organoid onto a 0.4-μm Transwell polyester membranes in 6-well plates (Corning). Following bioprinting, organoids were cultured for 1h in presence of 6μM CHIR99021 in TeSR-E6 medium in the basolateral compartment and subsequently cultured until day 12 in TeSR-E6 medium supplemented with 200 ng/ml FGF9 and 1 μg/ml Heparin. From day 12 to day 25, organoids were grown in TeSR-E6 medium either without additional supplement, or with additional 5uM all-trans retinoic acid (). Unless otherwise stated, kidney organoids were cultured until harvest at day 25.

### Flow cytometry

Prior to analysis, single kidney organoids were dissociated with 0.2 ml of a 1:1 TrypLE/Accutase solution in 1.5-ml tubes at 37°C for 15–25 min, with occasional mixing (flicking) until large clumps were no longer clearly visible. 1 ml of HBBS supplemented with 2% FBS was added to the cells before passing through a 40-lM FACS tube cell strainer (Falcon). Flow cytometry was performed using a LSRFortessa Cell Analyzer (BD Biosciences). Data acquisition and analysis were performed using FACSDiva (BD) and FlowLogic software (Inivai). Gating was performed on live cells based on forward and side-scatter analysis.

### Whole mount immunostaining

Fixed kidney organoids were incubated in blocking buffer (PBS 1X donkey serum 10% triton X100 0.3%) at 4°C for 3h before adding primary antibodies against HNF4α (Life Technologies 1:300, cat# MA1-199), Nephrin (NPHS1 1:300, Bioscientific, cat# AF4269) and Claudin-1 (CLDN1 1:100, Thermo Fisher Scientific, cat# 71-7800) at 4°C for 2 days. After washing in PBS 1X triton X-100 0.1%, organoids were incubated in secondary antibodies 1:400 at 4°C for 2 days: Alexa fluor 405 donkey anti-mouse (Abcam, cat# ab175659), Alexa fluor 488 donkey anti-goat (Molecular Probes, cat# A11055), Alexa fluor 568 donkey anti-rabbit (Life Technologies, cat# A10042). Samples were then washed before blocking at 4°C for 3h with PBS 1X mouse serum 10μg/ml triton X-100 0.3%, and adding an APC-conjugated CD31 antibody (1:50, Biolegend, cat# 303115) at 4°C for 2 days. Finally, samples were washed and imaged in 50:50 glycerol:PBS 1X using a Dragonfly Spinning Disc Confocal Microscope (Andor Technology).

### Single-cell transcriptional profiling and data analysis

Organoids were dissociated as described above (for flow cytometry) and passed through a 40-μM FACS tube cell strainer. Following centrifugation at 300 g for 3 min, the supernatant was discarded and cells resuspended in 50 μl TeSR-E6 medium. Viability and cell number were assessed, and samples were run across separate runs on a Chromium Chip Kit (10× Genomics). Libraries were prepared using Chromium Single Cell Li sequenced on an Illumina HiSeq with 100-bp paired-end reads. Cell Ranger (v1.3.1) was used to process and aggregate raw data from each of the samples returning a count matrix. Quality control and analysis was performed in *R* using the *Seurat* package (v3.2.2). Classification was performed using *DevKidCC* (v0.1.6) as described in this manuscript.

## Declarations

### Availability of data and materials

*DevKidCC* is available from Github at https://github.com/KidneyRegeneration/DevKidCC^64^ under the MIT licence. *DevKidCC Kidney Organoid Gene Expression* interactive shiny dashboard is available at https://sbwilson91.shinyapps.io/devkidcc_interactive/^59^ and from Github at https://github.com/KidneyRegeneration/DevKidCC_Interactive^59^

Single cell RNA-sequencing human fetal kidney datasets can be found in GEO (GSE102596, GSE114530) and EMBL-EBI ArrayExpress (E-MTAB-9083) ^60,62,65^. Single cell RNA-sequencing organoid datasets can be found in GEO (GSE118184, GSE109718, GSE119561, GSE114802, GSE115986, GSE132026, GSE124472, GSE152014, GSE161255, GSE152685)^61,66–73^. The single cell RNA-sequencing organoid dataset generated in this study will be available from GEO upon manuscript publication.

### Authors’ contributions

SBW, MHL and JEP conceived the study. SBW, JAH and JEP contributed to method development. SBW performed bioinformatics analysis. SBW, SEH, JMV and AD performed kidney differentiation experiments, immunofluorescence and FLOW analysis. SBW and MHL wrote the manuscript while all authors assisted in manuscript preparation.

## Acknowledgements

The authors would like to thank and acknowledge constructive discussions that contributed to this manuscripts development by all members of the Little laboratory and Powell laboratory

---

**Supplementary Figure 1:**
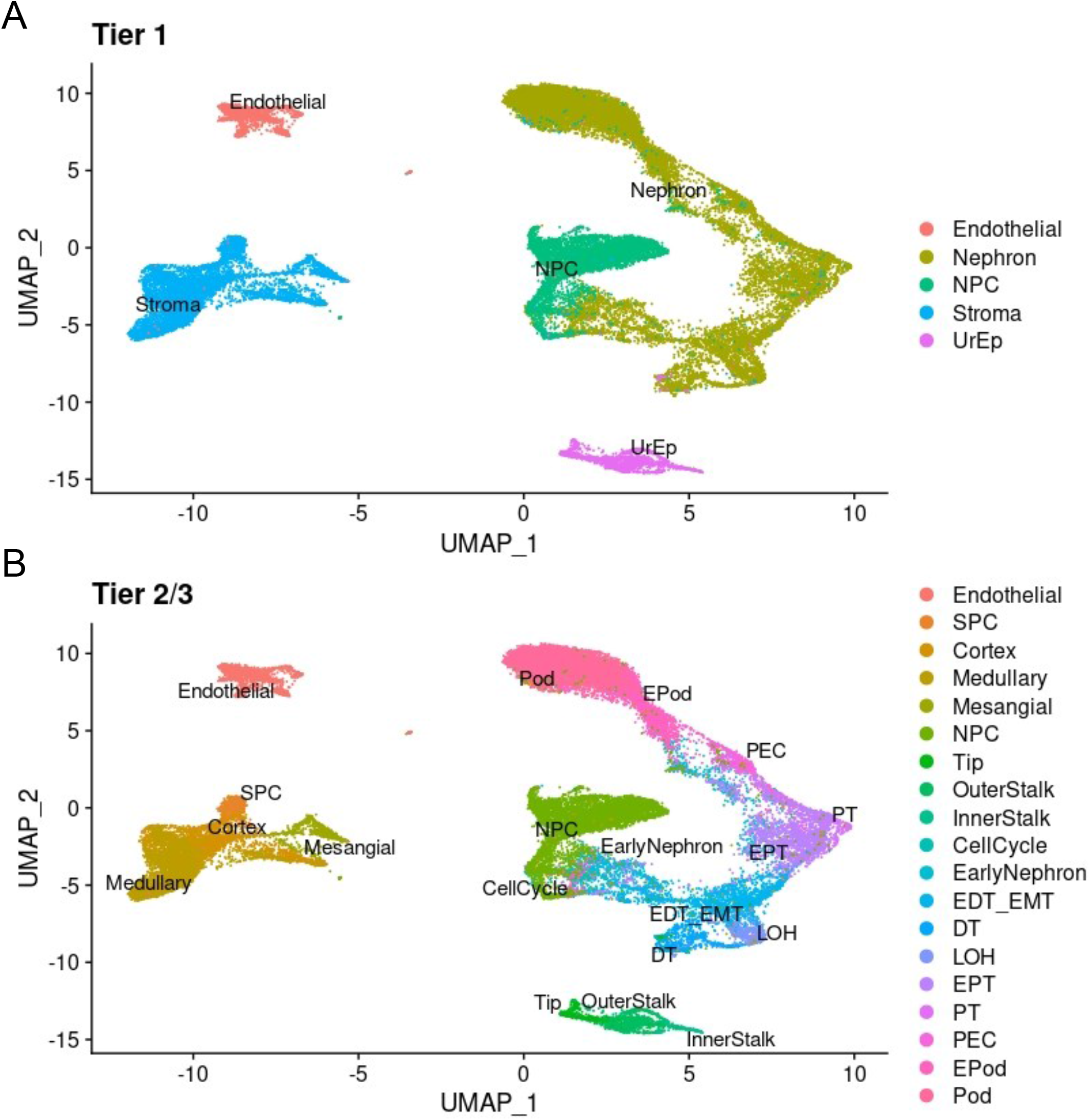
Annotation of the comprehensive reference. A) The reference annotation at Tier 1. B) The reference annotation at Tier 2.

**Supplementary Figure 2:**
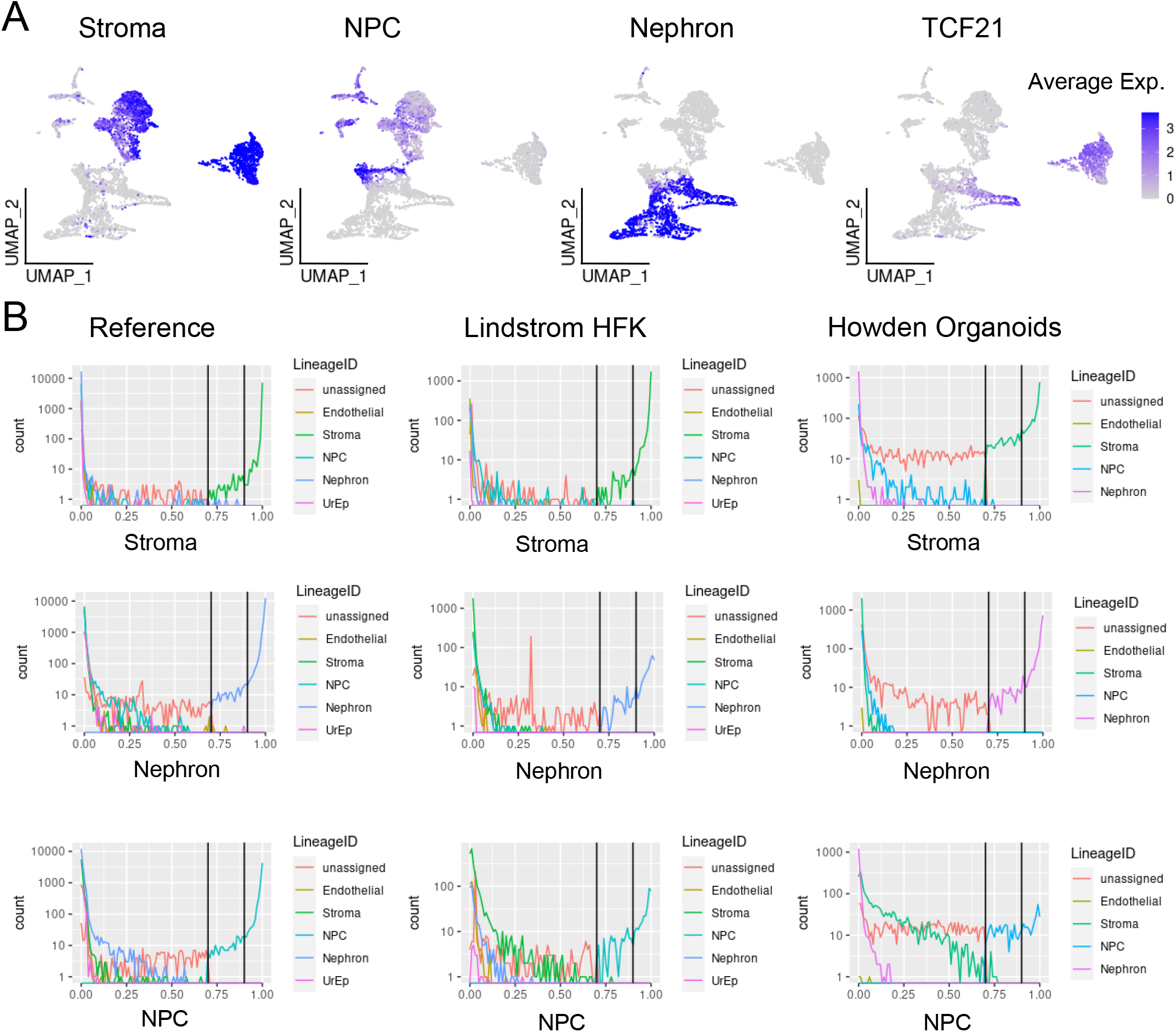
Scoring outcomes. A) UMAP plots showing the distribution of scores at the top tier for Stroma, NPC and Nephron, then the expression of *TCF21* which is kidney stromal marker. B) Density plots showing the distribution of cell scores for the Stroma, Nephron and NPC across the reference dataset, Lindstrom^19^ human fetal kidney (HFK) and Howden^13^ organoids datasets.

**Supplementary Figure 3:**
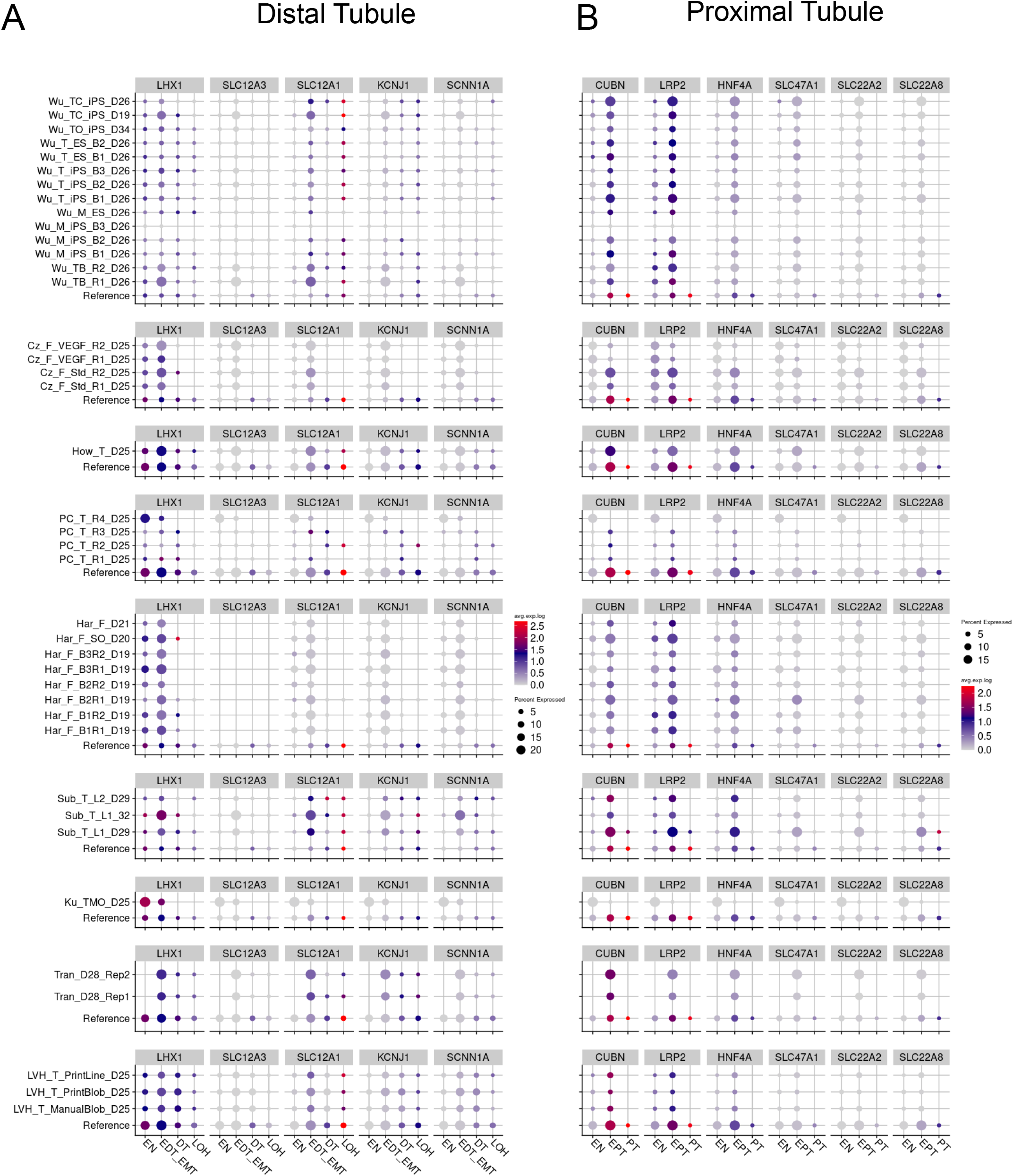
Epithelial maturation marker expression in end-stage organoids. Gene expression profiles for immature and mature markers of A) Distal Tubule and B) Proximal Tubule present in the relevant epithelial segments. *CUBN, LRP2, HNF4A, LHX1* are immature markers, *SLC47A1, SLC22A2, SLC22A8, SLC12A1, SLC12A3, KCNJ1, SCNN1A* are mature markers.

**Supplementary Figure 4:**
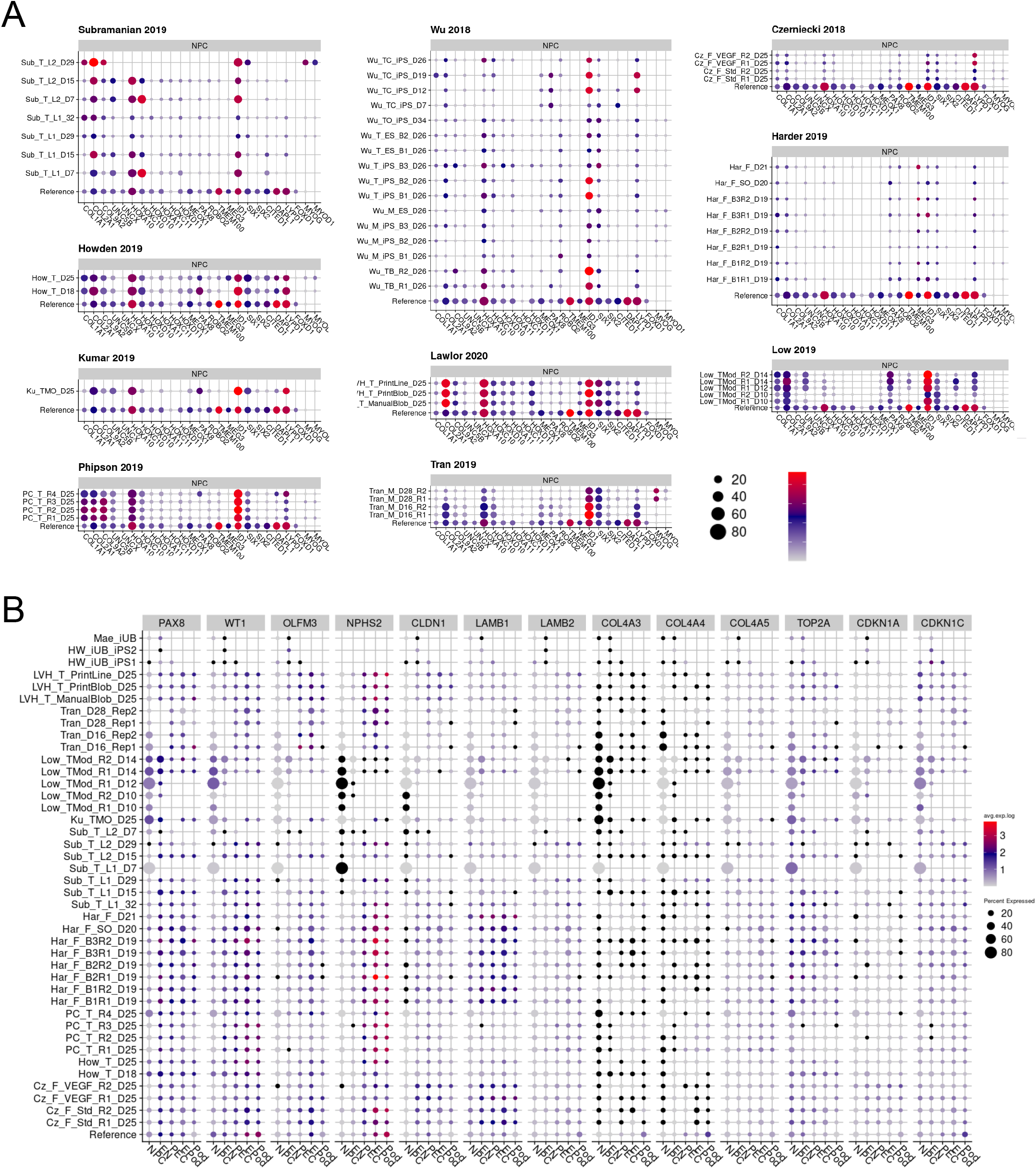
Expression of glomerular maturation markers in end-stage organoids. Gene expression profiles for markers of glomerular maturation. *WT1* is expressed from NPC through to mature podocytes. *OLFM3* is expressed in immature podocytes only. *NPHS2* is expressed in mature podocytes only. *CLDN1* is expressed in PECs. Immature podocytes express *LAMB1* and switch to *LAMB2* upon maturation. They also turn on *COL4A3*, *COL4A4* and *COL4A5* as they mature. Mature podocytes stop cycling and so are lowly expressing *TOP2A* and highly expressing *CDKN1A* (p21) and *CDKN1C* (p57).

